# Dehydrating microhabitats increase mite activity and intensify ectoparasitism of *Drosophila*

**DOI:** 10.1101/2025.09.24.678414

**Authors:** Joshua B. Benoit, Gabrielle LeFevre, Joy Bose, Ann Miller, David Lewis, Hailie Talbott, Chandrima Das, Emily Susanto, Lyn Wang, Oluwaseun M. Ajayi, Shyh-Chi Chen, Michal Polak

**Affiliations:** Department of Biological Sciences, University of Cincinnati, Cincinnati, OH 45211

**Keywords:** fly, mite, dehydration, parasitism, activity, resistance

## Abstract

Parasites interact with their host in variable environments that are often subject to water scarcity and dehydration. *Drosophila* fruit flies and associated ectoparasitic mites interact across a range of microhabitats, typically in decaying organic matter, such as fallen fruit and cactus tissue, that dries out and deteriorates over periods ranging from days to months. Here, we report that mite parasitism of *Drosophila* increases with exposure to increasingly dry conditions for two fly-mite systems (*D. nigrospiracula-Macrocheles* and *D. melanogaster-Gamasodes*). In *D. melanogaster*, artificial selection for behavioral resistance did not eliminate this effect, as previously selected lines remained relatively more resistant than non-selected controls even under dry conditions. Water balance assays confirmed that mites became dehydrated when held under dry conditions, which was also associated with increased mite activity. Exposure of *D. melanogaster* to mites dehydrated by exposure to low relative humidity increased parasitism, further supporting that mite infestation intensifies under dry conditions. The results indicate that ectoparasitism in this system is affected by the water content of the mites. The increased motivation of mites to parasitise flies under dry conditions may serve to replenish mite water stores and facilitate dispersal to more favorable microhabitats.

## Introduction

A diverse array of parasites exploit *Drosophila* fruit flies as hosts, and many of these associations are well-suited for experimental studies of host-parasite interactions. In particular, naturally occurring associations between *Drosophila* and ectoparasitic mites (Polak and Markow 1995; Halliday 2000; Halliday et al. 2005; Perez-Leanos et al. 2017; Yao et al. 2020) serve as tractable models for investigating the behavioral mechanisms insects use to resist infestation–mechanisms that are well characterized in vertebrate animals (Hart and Hart 2018). In general, mites are known to benefit from attaching to insect hosts through acquiring nutrients, increasing reproduction, and facilitating dispersal to new habitats (Walter and Proctor 2013). In *Drosophila*, mite attachment harms the host through direct damage to the cuticle and resource extraction, triggering wound repair, melanotic responses, and other immune responses (Benoit et al. 2020; Polak et al. 2025). The damage caused by parasitism can be severe with negative consequences for host reproduction and lifespan (Polak and Markow 1995; Polak 1996). Parasitism by mites is also associated with a suite of behavioral and physiological changes in the host, including upregulation of male reproductive effort (Polak and Starmer 1998), diminished copulation success owing to direct mechanical interference by attached mites (Polak et al. 2007), and increased metabolic rate (Brophy and Luong 2021).

*Drosophila melanogaster* and other fruit fly species are known to be parasitised by *Gamasodes* mites, with field records from Asia (Taiwan and Thailand) and Australia (Halliday et al. 2005; Yao et al. 2020; Polak 2025; Polak et al. 2025). In North America, the mite *Macrocheles subbadius* parasitizes *Drosophila nigrospiracula* in the Sonoran Desert (Polak and Markow 1995). This broad geographic distribution of fly–mite associations, together with their occurrence across diverse ecological settings, suggests that ectoparasitic mites exert widespread selective pressures on multiple aspects of fly biology. Flies avoid infestation by ectoparasitic mites by deploying rapid bursts of flight from the substrate, evasive locomotor maneuvers, and vigorous grooming once mite contact is made (Polak 2003; Polak et al. 2023). Behavioral resistance to mites is significantly heritable (Polak 2003; Luong and Polak 2007) and can be subject to strong selection (Polak and Markow 1995; Polak 1996; Benoit et al. 2025). Little is known about how environmental conditions experienced by mites and flies influence the outcome of these interactions.

Exposure to dry conditions and the prevention of dehydration are significant factors influencing the survival and proliferation of terrestrial arthropods (Benoit et al. 2023). When flies and mites interact, the water content of the local substrate, such as decaying fruits, flowers, cacti, or mushrooms, often declines, eventually reaching levels that cannot sustain many small arthropods, such as *Gamasodes* or *Macrocheles,* either due to this declining water content of the substrate or the availability of prey. These observations suggest that mites may increase their rates of parasitism on fruit flies when chronically exposed to dry off-host conditions, both to obtain water and nutrients from the fly and to disperse to more favorable microhabitats. For example, one previous study of the cactophilic fruit fly species *D. nigrospiracula*, found that both the intensity and prevalence of *M. subbadius* mite parasitism increased with the age and deterioration of necrotic saguaro cactus (*Carnegiea gigantea*) tissue, likely due to a combination of desiccation of the necrosis and declining habitat suitability for mite foraging and reproduction (Polak and Markow 1995). Several other studies suggest that ectoparasites and parasitoids increase their feeding and parasitism rates when exposed to dry conditions (Hagan et al. 2018; Bezerra Da Silva et al. 2019; Abu et al. 2024; Holmes et al. 2025). However, such interactions have yet to be directly explored in *Drosophila*–mite systems.

The present study investigated how dry conditions influence interactions between *Drosophila* fruit flies and their ectoparasitic mites. Our experiments, which examined two distinct fly–mite systems, *D. nigrospiracula-Macrocheles* and *D. melanogaster-Gamasodes*, demonstrated that dehydration increases mite parasitism in both systems. In *D. melanogaster*, flies that were previously artificially selected for parasite resistance (Polak et al. 2023; Benoit et al. 2025) exhibited greater resistance to mite attachment than non-selected flies under both dry and moist conditions, and both groups became more heavily parasitized when mites had been exposed to dry conditions. Thus, dry conditions overall significantly increased parasitism rate. Water balance assays revealed that mites held under dry conditions exhibited reduced water content, which correlated with heightened mite activity levels. Importantly, exposure of *D. melanogaster* to dehydrated mites resulted in increased parasitism of flies, supporting a link between substrate quality, mite physiological state, and parasitism. Our findings suggest that dry conditions directly increase mite motivation to parasitize fruit flies, which are likely to obtain water and nutrients from their hosts and be carried to more favorable microhabitats.

## Materials and methods

### Flies, mites and selection for mite resistance

A population of *Drosophila melanogaster* Meigen (Canton-S, number 64349) was obtained from the Bloomington *Drosophila* Stock Center and used in all experiments with this species described here, except those involving lines artificially selected for mite resistance. The selected lines were derived from a single outbred base population established with field-collected flies from Cape Tribulation, Australia (Polak et al. 2023). Artificial selection for increasing resistance, described in detail elsewhere (Polak 2003; Polak et al. 2023), was conducted using mites from a laboratory culture established with individuals harvested from flied-caught flies at Cape Tribulation (Polak et al. 2023). The selection protocol generated three replicate lines of *D. melanogaster*, each showing significantly greater resistance than its paired unselected control line (Polak et al. 2023). For the present study, flies were used after 22–24 generations of selection. Cultures of *Drosophila nigrospiracula* Patterson and Wheeler and their ectoparasitic mites *Macrocheles subbadius* Berlese were established from collections on *Carnegiea gigantea* cacti near Phoenix, Arizona, USA.

*Drosophila melanogaster* was cultured in 4 half-pint glass bottles per generation containing a cornmeal-agar food substrate (Webster and Polak 2025), while *D. nigrospiracula* was cultured in bottles with a food substrate composed of powdered potato flakes, instant *Drosophila* medium, and necrotic cactus (Polak 1998). All fly cultures were maintained in an environmental chamber under standard 12h light (24°C):12h dark (22°C) conditions. Both mite species were cultured in 5 L plastic jugs with a specialized medium (Polak 2003), and maintained separately in environmental chambers at 12h light (25°C):12h dark (24°C) conditions. The culture medium for both mite species consisted of a rich organic substrate of autoclaved wheat bran and wood shavings, supplemented with autolyzed yeast and bacteriophagic nematodes as a food source for the mites (Polak 2003; Benoit et al. 2020; Polak et al. 2023). The culture medium was moistened with sufficient deionized water to keep it damp without puddling. Dry stock medium used in the experiments described below consisted of autoclaved wheat bran, wood shavings, and autolyzed yeast. Male flies were used at 7-10 days of age.

### Dehydration treatment

To generate “dry” and “wet” culture conditions, a portion of the mite-containing culture medium was replaced with either dry or hydrated stock medium. This was accomplished by first removing ∼30-40% of the culture medium and replacing it with stock medium that was either dry or wet. The media were mixed by gently shaking the container for one minute until the mixture was homogeneous. After 24 hours, the medium was observed for the presence of mites. If mites were present, medium and mites were collected to determine their water contents. Rehydrated medium was generated for a subset of the dry treatments by adding water to the container, allowing the mites to rehydrate for 24 hours before conducting mite infestation assays.

### Media water content determination

A standard soil moisture content assay (Basimike and Mutinga, 1990; Su and Puche, 2003) was used to determine the proportional change in water content of the mite media as a result of making them either drier or wetter. One day after the addition of dry or wet media, the top 2 cm was removed and immediately weighed on aluminum pans. Following the determination of the initial mass (wet mass), the media were dried at 0% RH and 50°C within a drying oven for two weeks. Following drying, they were weighed on three consecutive days to determine that the water lost from the soil had reached the dry mass (no additional decline in mass). Water content was determined as a percentage of the initial mass.

### Mite infestation experiments

Mite infestation assays followed established methods to test for differences in susceptibility between experimental groups of flies (Polak 1996, 2003; Luong and Polak 2007; Benoit et al. 2020). Briefly, flies were placed in infestation chambers (300 mL glass jars) containing culture medium with mites (Polak 1996, 2003; Benoit et al. 2020), similar to the chambers used during artificial selection for increased resistance in *D. melanogaster* (Polak et al. 2023). For both *D. melanogaster* and *D. nigrospiracula*, groups of 40-50 flies were added to each infestation chamber; *D. melanogaster* was exposed to *G. queenslandicus* mites, whereas *D. nigrospiracula* was exposed to *M. subbadius* mites. Only male flies were used in the study.

To assess the effect of the moisture content of the substrate on infestation, assays compared ectoparasitism rates under dry versus wet conditions across replicate infestation chambers for each host species. Infestation chambers containing dry or wet media were run in parallel to ensure exposure times were similar between treatments. Approximately the same volume of media was added to the chambers to help maintain similar numbers of mites. After exposure, flies were extracted from chambers with an aspirator, and under CO₂, the presence of mites and scars was assessed for each fly. Prevalence of parasitism ((scarred plus parasitized flies)/total flies exposed) was calculated for each group.

For the *D. melanogaster* selection lines, two groups of flies were exposed to mites in a common infestation chamber, one group from a given selected line and other from its paired non-selected line. Group sizes were approximately equal, with a total of 40–50 flies per chamber. To distinguish the two groups, a minute wing clip was administered to the tip of either the right or left wing (≤ 3% of wing area) while flies were under CO_2_. Prior to aspirating them into chambers, flies were allowed to recover from the CO_2_ for 24 hours. Wing clips have been previously shown to have no significant effect on fly susceptibility to mites (Polak 2003; Benoit et al. 2020). The wing that received a clip was alternated between groups across chambers. Flies were aspirated into infestation chambers and allowed to interact with mites for 6–8 hours in darkness, depending on mite density in the medium. After exposure, flies were extracted from chambers with an aspirator, and under CO₂ anesthesia, their identities ascertained by wing clips, and the prevalence of parasitism determined as described above.

In separate assays, mites were exposed to dehydrating conditions in the absence of culture medium. Briefly, mites were removed from the media and individually placed in mesh-covered 0.5 ml microcentrifuge tubes. The tubes were placed at either 33% relative humidity (RH, maintained with saturated MgCl_2_ solution in water) or 93% RH (maintained with saturated KNO_3_ solution in water) in glass desiccators for 6-8 hours before exposing the flies to the mites. One fly and one mite were put together in a standard *Drosophila* vial (25 x 95 mm), which was then kept in a desiccator at 93% RH. Incidence of parasitism was evaluated after 30 minutes.

### Mite water content determination

Mite water content was determined using protocols previously applied to other mite species (Benoit et al. 2008; Yoder et al. 2012). Mites (deutonymphs) were collected directly from the media, and their initial mass was determined within five minutes after freezing at −20°C. A Cahn microbalance was used to measure the mass of individual mites to the nearest 0.1 μg. Dry mass was determined by drying mites at 0% relative humidity (using CaSO₄, Drierite) at 50°C until stable weights were obtained over three consecutive days; the final value was taken as the dry mass. Water content was defined as the difference between wet and dry mass.

### Activity assays

Mite activity was assessed following methods developed for small arthropods (Fieler et al. 2021; Bailey et al. 2024; Benoit et al. 2025). As before, mites were individually placed in mesh-covered 0.5 ml microcentrifuge tubes and held either at 75% RH or 93% RH for 6 hours. Groups of three mites were collected and transferred to 10-cm glass tubes, which were sealed at both ends with foam stoppers. Tubes were placed horizontally into a locomotor activity monitor (LAM, Trikinetics). Groups of three mites, rather than individuals, were used to assess activity to ensure that enough mites passed through the infrared beams that serve to detect movement (Fieler et al. 2021; Benoit et al. 2025). The activity monitor was placed in darkness at 95-98% RH within a Percival incubator. After a two-hour acclimation period, mite behavior was recorded for six hours. Activity was quantified as the number of beam crossings per hour. To minimize circadian effects, measurements were taken nightly from 8 PM to 8 AM, as mites show marked differences in activity between day and night (Benoit et al. 2025).

### Statistical analyses

R software (Version 4.3.1) was used for statistical analyses and figure development. General linear models (binomial for proportion and gamma for non-proportion data) were used to examine statistical differences between treatments (dry vs. wet conditions) for each species. A Generalized Linear Mixed Model (GLMM - binomial) was used to compare the results of selection lines in relation to hydration status. In the models, the infestation chamber was added as a random effect to account for differences among chambers, due, for example, to mite density variation. In all cases, chamber effects were not significant (P > 0.2 for all models). Statistical outputs and sample sizes are provided within the figure legends.

## Results

### Dry conditions increase mite parasitism in two fly-mite systems

When we examined the water content of wet vs. dry media, there was a significant 30-40% decline in the water content of the media (Fig. 1). When *D. melanogaster* was exposed to *G. queenslandicus*, a little over half of the flies became parasitized with mites (Fig. 2A). Dehydrating conditions resulted in a significant increase in parasitism, with 25-30% more flies harboring mites (Fig. 2A, P < 0.0001). A similar effect of medium dehydration was noted for *D. nigrospiracula* interacting with *M. subbadius* mites (Fig. 2B; P = 0.0002), but the only difference was that overall parasite levels were lower compared to *D. melanogaster*–*G. queenslandicus* interactions. The addition of water to the dry media resulted in a reduction in mite parasitism for *D. melanogaster* (Fig. 3, P < 0.0005), confirming that dry conditions contributed to increased mite infestation of the flies.

**Figure 1.**
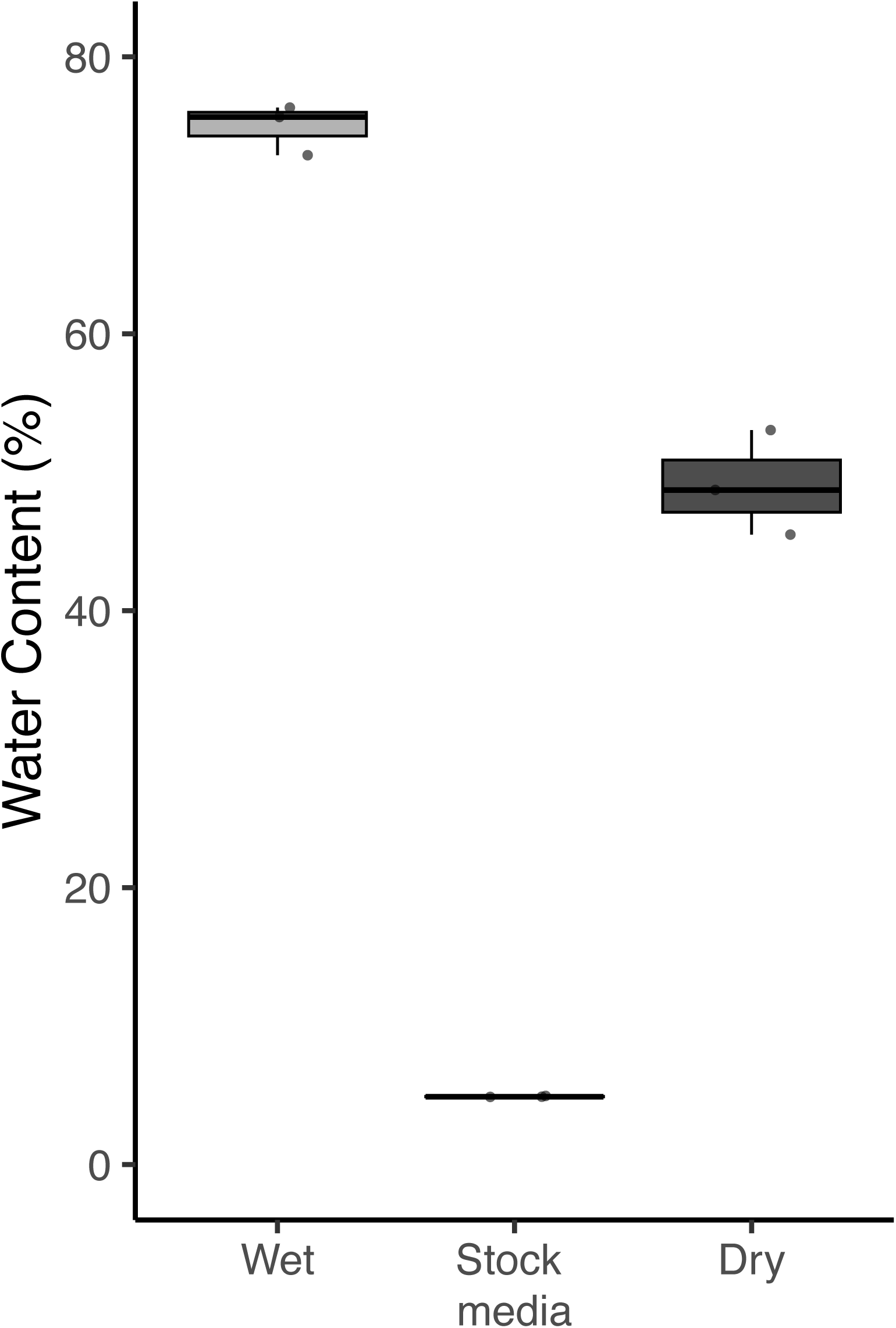
Water content of mite media under wet and dry conditions. Water content of mite media in dry or wet conditions, along with the water content of the dry stock medium (N = 3 samples for each media type). F_2,6_ = 648.5, P < 0.0001, all media types are significantly different (P < 0.005).

**Figure 2.**
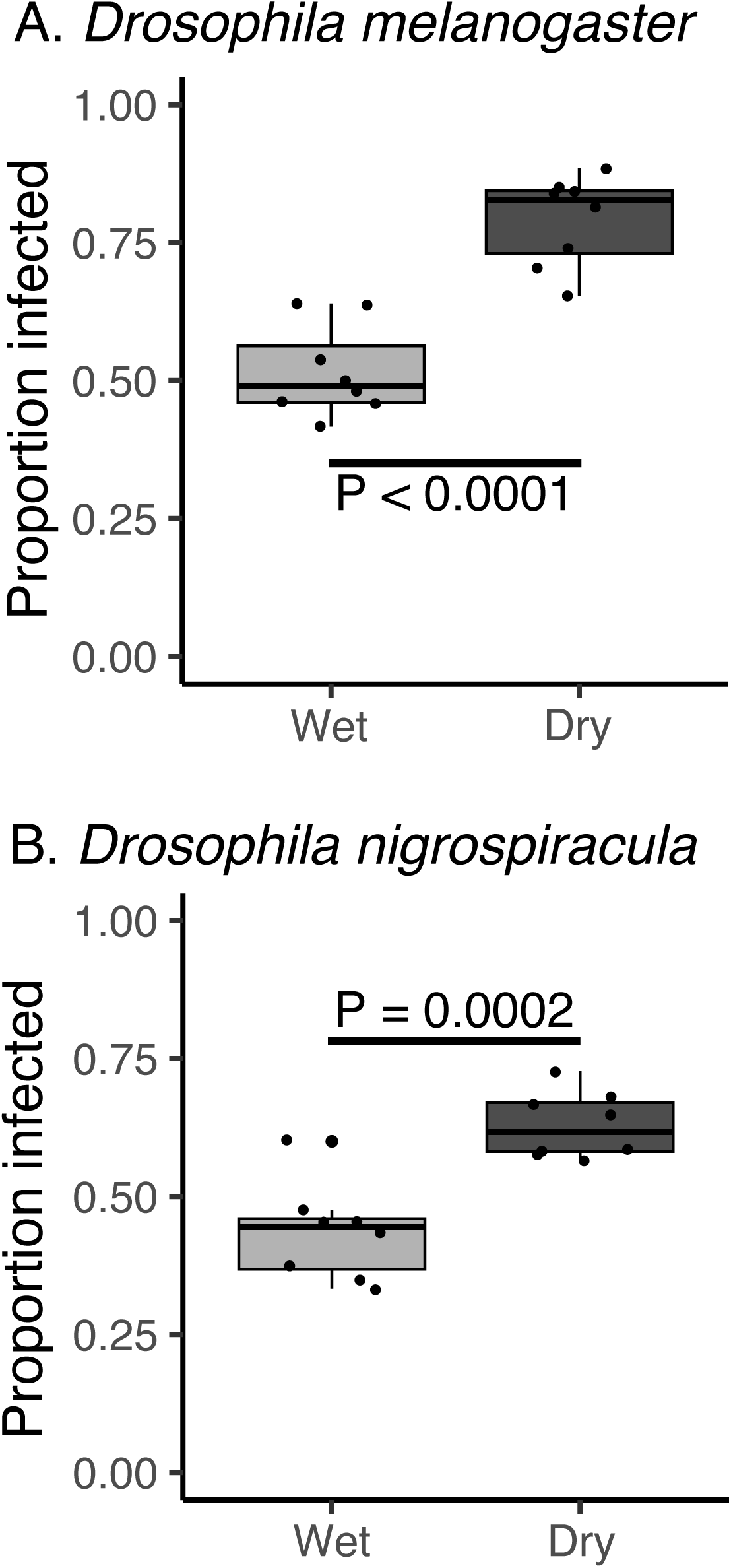
Dehydration increases mite parasitism on fruit flies. A. *Drosophila melanogaster* (Canton S) exposed to mites (*Gamasodes queenslandicus*) under wet and dry conditions. N = 8 chambers per condition, P < 0.0001. B. *Drosophila nigrospiracula* exposed to mites (*Macrocheles subbadius*) under wet and dry conditions. N = 8 chambers per condition, P = 0.0002. There was a general effect where both species showed increased parasitism under dry conditions, F_1,30_ = 43.64, P < 0.0001.

**Figure 3.**
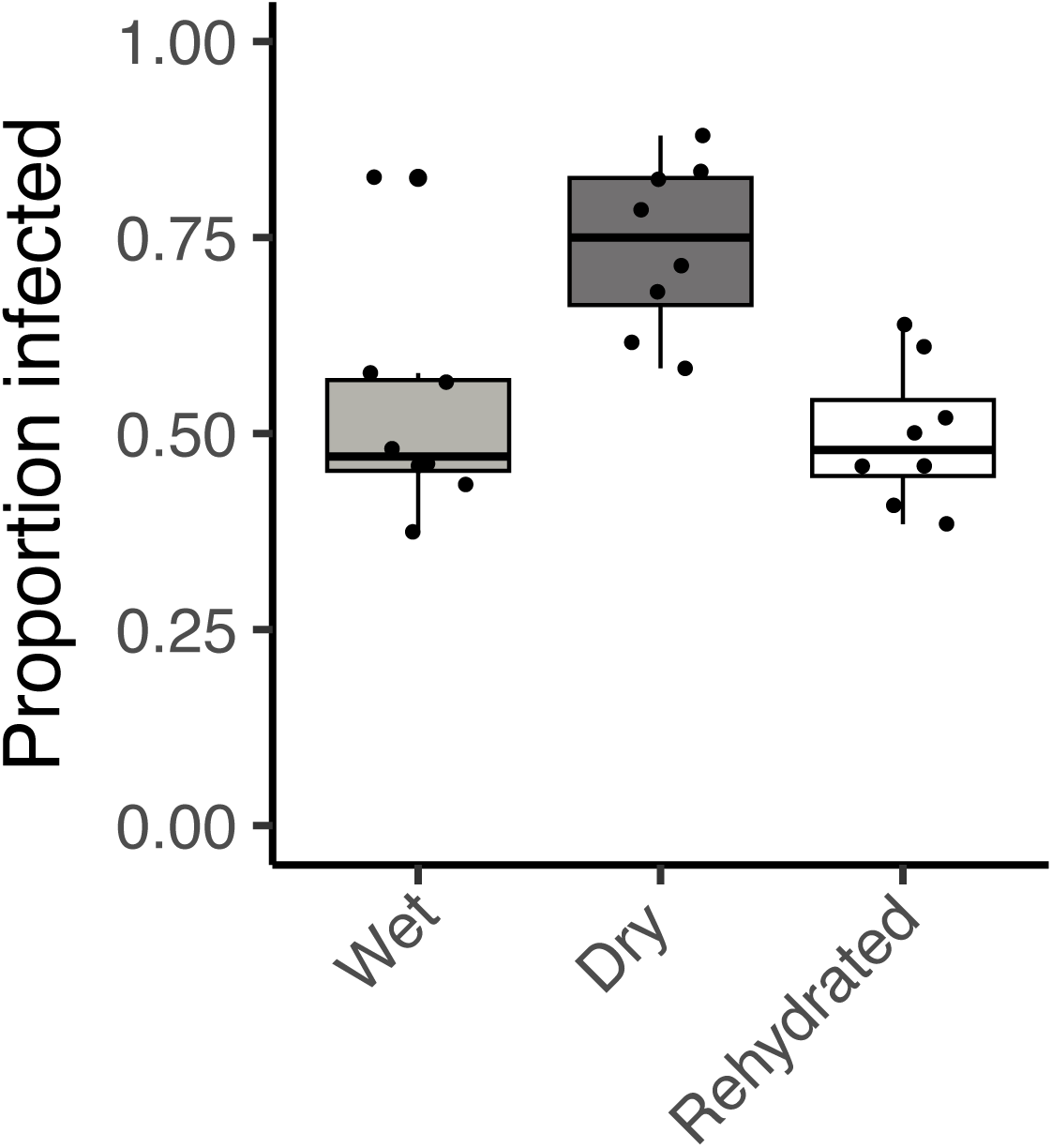
Rehydration of media reduces parasitism to levels in wet media. *Drosophila melanogaster* (Canton S) exposed to mites (*Gamasodes queenslandicus*) under wet and dry conditions, along with those exposed to dry medium that was rehydrated for 24 hours. N = 8 chambers per treatment. An increase in parasitism was noted only under dry conditions compared to both wet and rehydrated conditions, F_2,21_ = 10.81, P < 0.0005.

### Selection for behavioral resistance functions under dry conditions

A previous study performed artificial selection for increased behavioral resistance of *D. melanogaster* against *G. queenslandicus* mites (Polak et al. 2023; Benoit et al. 2025), generating the selected and non-selected lines used in the present study. When selected and non-selected flies were exposed to mites in the same chamber, dry conditions increased parasitism in both groups compared to wet conditions (Fig. 4, P < 0.005). Selected flies in both environments maintained their resistance superiority, with non-selected flies exhibiting 12.8% and 11.8% higher parasitism under wet and dry conditions, respectively (Fig. 4).

**Figure 4.**
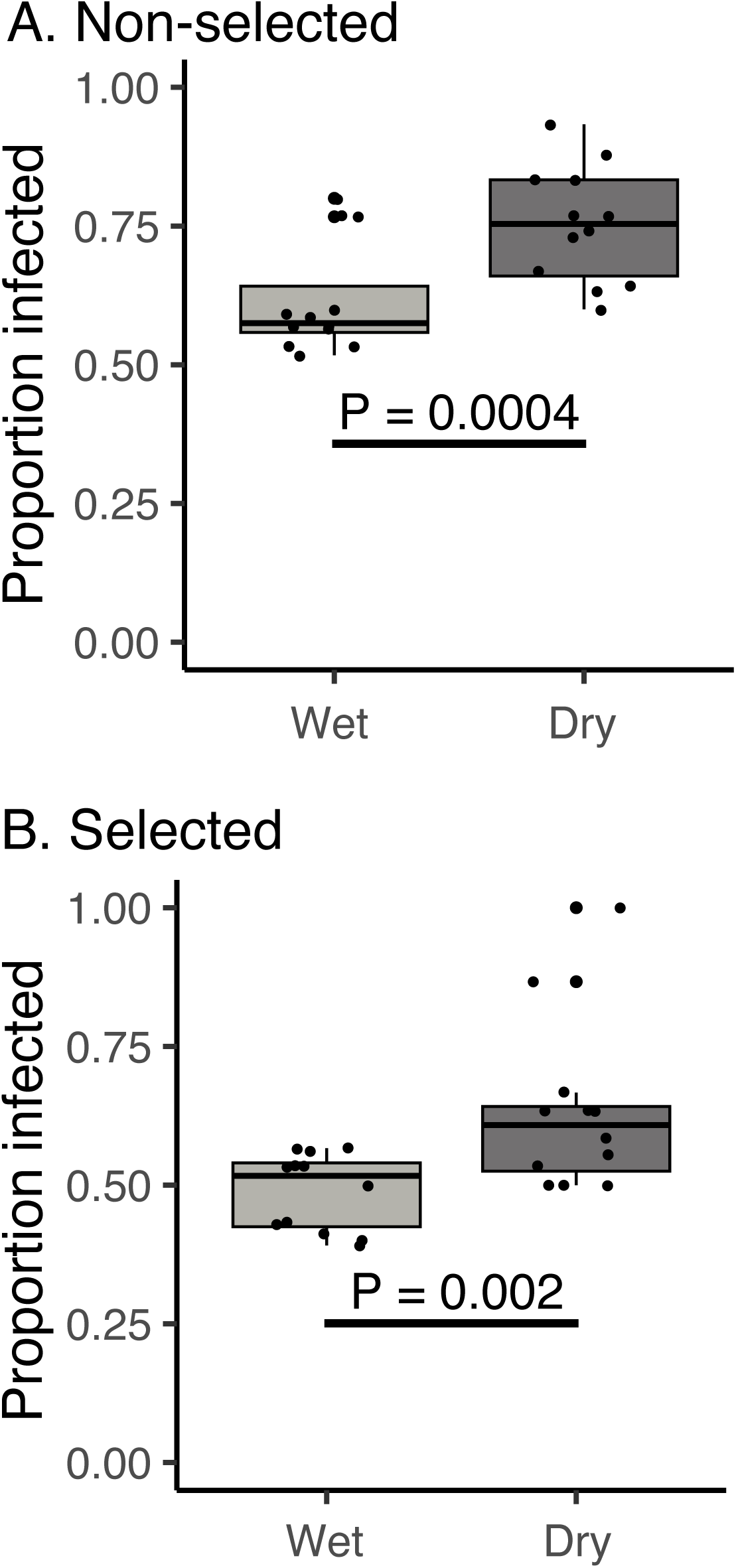
Dehydration increases mite parasitism for mite-resistant and non-resistant lines. Both non-selected (A) and selected (B) lines were parasitized at higher rates under dry conditions compared to wet conditions (P < 0.002 for both cases). Selected lines were more resistant than non-selected lines under both wet (P=0.001) and dry (P=0.003) conditions. N = 12 chambers per treatment.

### Dry conditions lead to a decrease in water content in mites

We evaluated whether the water content of *Gamasodes* mites differed between wet and dry conditions (Fig. 5). Mites exposed to dry conditions within infestation chambers had significantly lower initial (wet) mass compared to those sampled from wet conditions (Fig. 5A; P = 0.002), but dry mass did not differ between the two groups (Fig. 5B; P = 0.374). Mites from the dry medium had a 20–25% reduction in water content relative to those from the wet medium (Fig. 5C; P = 0.0003). This water balance assessment confirms that *G. queenslandicus* mites become dehydrated when exposed to dry conditions.

**Figure 5.**
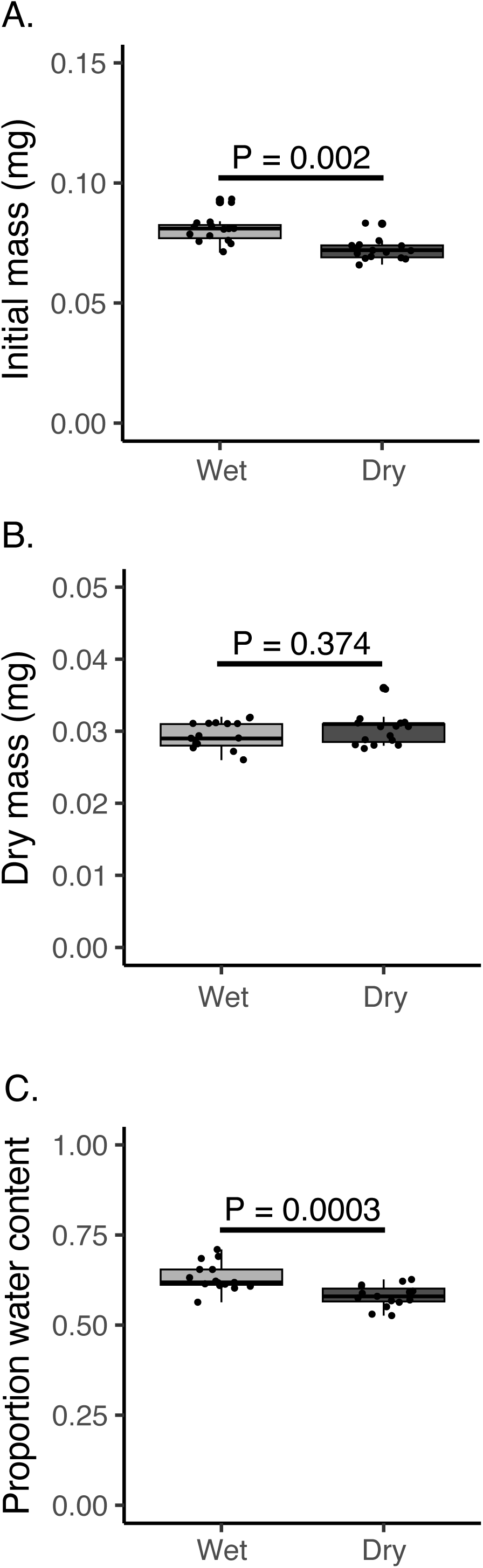
Water content mites from wet and dry media conditions. A. Initial mass. F_1,28_ = 23.26, p = P < 0.001. B. Dry mass. F_1,28_ = 0.8178, P = 0.0374. C. Water content of mites. F_1,28_ = 17.19, P = 0.001. N = 15 for mites held in wet and dry media.

### Dehydrated mites exhibit increased activity and are more likely to parasitize flies

Dehydrated mites exhibited significantly higher activity levels than those from moist media, showing a 20–30% increase in activity (Fig. 6A, P = 0.006). When mites were exposed to the flies in a media-free chamber, the dehydrated mites were nearly 2-fold more likely to parasitize the flies compared to hydrated mites (Fig. 6B, P = 0.012).

**Figure 6.**
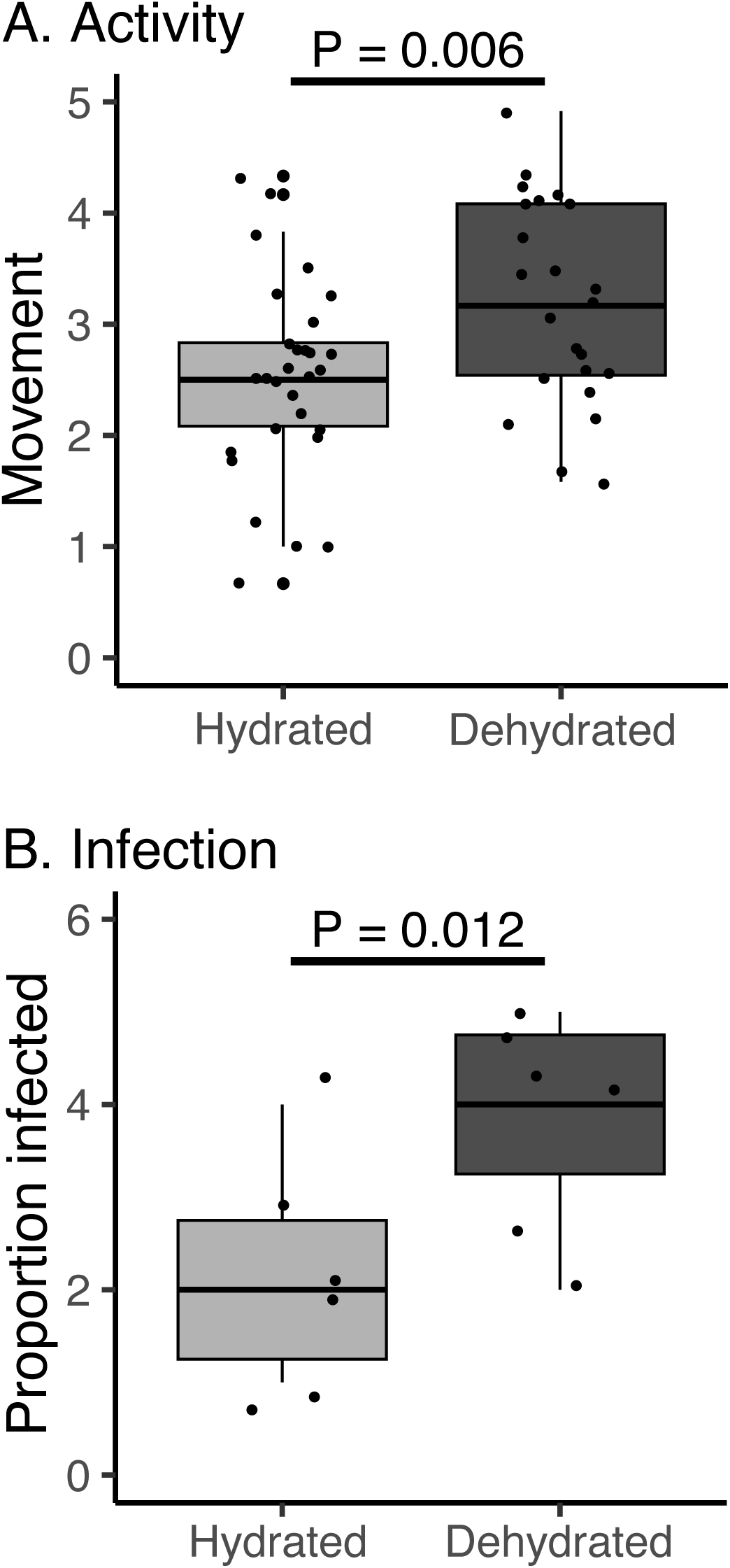
Activity levels and parasitism are increased in dehydrated mites. A. Activity levels were higher in dehydrated mites compared to hydrated mites, N = 30 for hydrated and N =28 for dehydrated mites. F_1,56_ = 8.01, p = 0.006. B. Direct assessment of mite-fly interactions confirms that dehydration triggers increased parasitism. N = 6 exposures of flies to the mites for hydrated and dehydration treatments. F_1,10_ = 9.31, p = 0.012.

## Discussion

This study demonstrated that dry substrate conditions are a significant factor driving parasitism by mites. In two independent fly-mite systems, dehydration significantly increased rates of parasitism. Interestingly, even after artificial selection for resistance in *D. melanogaster* (Polak et al., 2023a), desiccation still enhanced parasitism, although selected flies retained elevated resistance when compared to their non-selected counterparts. Water balance measurements in *G. queenslandicus* mites confirmed reduced body water content under dry conditions. Dehydrated mites also exhibited increased locomotor activity, which likely contributed to heightened host encounter and attachment rates. However, this increased activity is perhaps best interpreted as part of a broader suite of mite behaviors. These include heightened host-seeking, which directly contributes to higher parasitism rates, as well as dispersal efforts to escape desiccating conditions through independent movement to wetter microhabitats. Together, these findings indicate that desiccating conditions can intensify host–parasite interactions by affecting off-host parasite behavior.

The impact of dry conditions on arthropod pests, especially ectoparasites has been a recent focus of several studies. In mosquitoes and ticks, dry conditions increase activity, host seeking, and feeding (Rosendale et al. 2016, 2019; Hagan et al. 2018; Manzano-Alvarez et al. 2023; Abu et al. 2024), as well as even drive male mosquitoes to ingest blood (Bozic et al. 2024). In addition to increased feeding, ectoparasites often exhibit behavioral shifts in response to dehydration, including elevated general activity and heightened movement after a bloodmeal (Rosendale et al., 2016; Holmes et al., 2025). Similar patterns have been observed beyond vertebrate–arthropod systems; for instance, water deprivation in insects has been linked to increased parasitism by parasitoid wasps (Bezerra Da Silva et al., 2019). Our results align with these previous studies, showing that two species of ectoparasitic mites that naturally parasitize fruit flies exhibited increased rates of parasitism under dry conditions that demonstrably promote mite dehydration. These results support the hypothesis that dehydration is a general environmental factor capable of intensifying ectoparasitism. Importantly, the observed increase in ectoparasitism is unlikely to be the result of mite starvation. The exposure period was relatively short, and prior work indicates that more extended deprivation is required to induce increased feeding due to starvation (Luong et al. 2017). Additionally, mites likely retained access to nematode prey within the dry and wet medium, as fresh media was added to both for only a single day. Thus, the increase in parasitism may more plausibly be attributed to dehydration-induced increases in mite activity, which likely enhanced host encounter rates. Further work will be required to distinguish the relative importance of dehydration versus starvation in driving elevated parasitism.

Resistance to mites confers fitness benefits to flies by reducing the probability of infestation (Luong and Polak 2007; Polak et al. 2023; Benoit et al. 2025). However, resistance is not cost-free: in the absence of parasites, elevated resistance may incur fitness costs, such as reduced stress tolerance and diminished fecundity, as documented in this and other host–parasite systems (Ewald 1995; Windsor 1998; Fitze et al. 2004; Luong and Polak 2007). These costs are usually more muted than the direct fitness consequences of parasitism, which can lead to strong physiological declines and increased probability of death (Polak and Markow 1995; Benoit et al. 2020; Polak et al. 2023). However, little is known about how the outcomes of host-ectoparasite interactions shift under suboptimal conditions. In the present study, we demonstrated that dry conditions elevated infestation rates in both selected and non-selected flies, indicating an effect of substrate humidity on the prevalence of parasitism. Interestingly, selected lines retained an advantage over their non-selected counterparts under both wet and dry conditions, indicating stability of the genetic effect on behavioral immunity across these environmental conditions.

The increased parasitism by mites when held under dry conditions we documented could represent a critical mechanism for mite dispersal to more optimal habitats. Flies utilize a wide range of substrates, ranging from fermenting fruit, cactus, and fungi, among other organic materials (Markow and O’Grady 2006). As these organic substrates are consumed, decompose, and become dry, particularly in areas of seasonal periods of limited rainfall, they not only become increasingly suboptimal for fly survival and reproduction, but these environmental effects are also felt by the mites themselves. Depletion of available nematodes and other small invertebrate prey for the mites may lead to mite starvation, which has previously been shown to increase mite motivation to parasitise flies (Luong et al., 2017). Furthermore, egg viability is expected to decline under desiccating conditions, a pattern commonly observed in mite eggs exposed to dry environments (Benoit et al. 2007; Yoder et al. 2012), thus reducing mite survival in microhabitats that dry as they deteriorate. Given that mites commonly parasitize flies for many hours (Webster and Polak 2025), and that parasitism by mites improves their fecundity (Luong and Subasinghe 2017), movement with the flies likely enhances mite fecundity and facilitates locating new, more optimal habitats (Polak and Markow 1995).

## Acknowledgments

The research was supported by National Science Foundation (NSF) grant 1654417 (to M.P. and J.B.B.). Partial funding for reusable equipment was provided indirectly by the National Institute of Allergy and Infectious Diseases, award numbers R01AI148551 and R21AI166633 to J.B.B.

